# Density-dependent feedback limits the spread of a beta-lactamase mutant: experimental observations and population dynamic model

**DOI:** 10.1101/2023.09.02.555988

**Authors:** Philip Ruelens, Eline de Ridder, J. Arjan G.M. de Visser, Meike Wortel

## Abstract

β-lactamases play an important role in antibiotic resistant bacterial infections. Understanding the spread of these enzymes may inform the development of better drug therapies. However, this is complicated by the fact that β-lactamases reduce the antibiotic concentration in their environment, thereby altering their own selective advantage via eco-evolutionary feedback. We investigated the effect of such feedback on the spread of bacterial strains expressing β-lactamase enzymes conferring different levels of resistance to the cephalosporin cefotaxime. Specifically, we conducted head-to-head competitions between two clinically observed β-lactamase mutants, TEM-19 and TEM-52, with low and high activity against cefotaxime, respectively. By experimentally varying nutrient levels, we altered cell density and hence the strength of ecological feedback and examined its impact on competitive fitness and strain coexistence across a range of cefotaxime concentrations. A population dynamic model, parameterized solely with independently measured traits, revealed cell density as the key mediator of this feedback. Our results show that cell density dictates whether the resistant strain (TEM-52) outcompetes or stably coexists with the susceptible strain (TEM-19). By validating our model with experimental data, we showed that it can reasonably predict the equilibrium frequencies based on dose-dependent growth rates and antibiotic degradation rates of both strains. Our study emphasizes the importance of considering ecological feedback for understanding the fate of antibiotic-degrading mutants, including in clinical environments.

**Importance:** Since the discovery of penicillin, β-lactam antibiotics have become the most widely used antibiotics to treat bacterial infections. Their applicability is decreasing because bacteria evolve resistance via expression of antibiotic-degrading β-lactamase enzymes. Because the spread of resistance is a large health problem, th prediction of resistance evolution is a big target. Several mutations increasing resistance have been studied, but as enzymes with increased activity spread, cells deactivate the β-lactam antibiotics and thereby open a niche for susceptible variants. This eco-evolutionary feedback complicates the prediction of resistance evolution and makes it context dependent. Here, we show that both the cell density and the antibiotic concentration affect the success of a new β-lactamase variant and whether it can invade, replace, or coexist with an ancestor variant. With a mathematical model we can predict the success of new variants and therefore predict evolutionary paths depending on the environmental variables.

## Introduction

Antibiotic resistance is a major global health threat that poses a significant challenge to modern medicine (1). The emergence and spread of resistant bacterial strains have rendered many antibiotics ineffective, leading to longer hospital stays, and an increase in morbidity and mortality (2). β-lactamases, a family of enzymes that hydrolyze β-lactam antibiotics, are among the most common and clinically relevant antibiotic resistance determinants (3). Given their importance, the evolution of β-lactamase resistance has been extensively studied (4–6). A factor complicating the evolutionary success of novel β-lactamases, that has been ignored in most studies, is the influence of eco-evolutionary feedback due to antibiotic removal by the β-lactamases themselves on the invasion of new β-lactamase variants (7–12). A more quantitative understanding of the effect of eco-evolutionary feedback on the spread of β-lactamases is required for a comprehensive understanding of their contribution to antibiotic resistance.

In population dynamics, invasion fitness is a common metric to assess whether a mutant can successfully invade and establish itself within a population (13). Negative frequency-dependent selection, where the benefit of a mutant decreases when it becomes more frequent, can prevent fixation and cause coexistence of more and less resistant genotypes. In the context of antibiotic resistance, this can occur when resistance mutations decrease the environmental antibiotic concentrations, thereby benefitting less resistant mutants — a phenomenon known as cross-protection (7,14–17). Cross-protection has been observed in high-density bacterial populations (7,11,14,17–19), including in clonal populations, where the effectiveness of an antimicrobial agent may decrease with increasing bacterial density, referred to as the ‘inoculum effect’ (18). Opportunities for the cross-protection of a susceptible strain by a β-lactamase producing resistant strain generally depend on the antibiotic concentration together with the initial cell density, dose-dependent fitness and antibiotic-degrading capacity of both strains, leading to a ‘cross-protection window’ (17).

For cross-protection to lead to coexistence, it is necessary that resistance confers a fitness cost – the susceptible strain needs to grow faster without antibiotics to compensate for its growth inhibition in the presence of antibiotics. This cost is often evident when antibiotic resistance determinants are located on a plasmid, but it less clear and can be variable for single nucleotide polymorphisms affecting the activity of a β-lactamase (20). As a result, the extent to which competing bacteria expressing different β-lactamases may influence each other’s spread through eco-evolutionary feedback remains unclear.

In order to successfully spread, bacteria expressing a novel β-lactamase with higher hydrolyzing capacity must outcompete the bacteria from which they derive. Essentially, both strains engage in a competition for limited resources, while particularly the more resistant strain removes antibiotic from the environment through enzymatic degradation (Figure 1). When the initial concentration of the antibiotic is above the minimum inhibitory concentration (MIC) of the more susceptible strain, the resistant strain may rescue the susceptible strain if the antibiotic concentration is reduced below the MIC of the susceptible strain before it is eliminated by the antibiotic. If so, the susceptible strain can resume growth and increase in frequency if the resistant strain has a lower fitness in the absence of antibiotic (18). Population density plays a crucial role in these competitive interactions, as high-density bacterial populations can degrade antibiotics more rapidly, enhancing the probability of susceptible bacteria to survive until the antibiotic concentration decreases below inhibitory concentrations. At low densities, in contrast, resistant bacteria can grow longer without competition from susceptible bacteria, allowing their relative frequency to increase more or the susceptible population to be killed entirely. Notably, when the resistant strain has lower fitness than the susceptible strain in the absence of antibiotic, conditions allowing the rescue of the susceptible bacteria by the resistant strain may lead to the stable coexistence of both strains.

**Figure 1.**
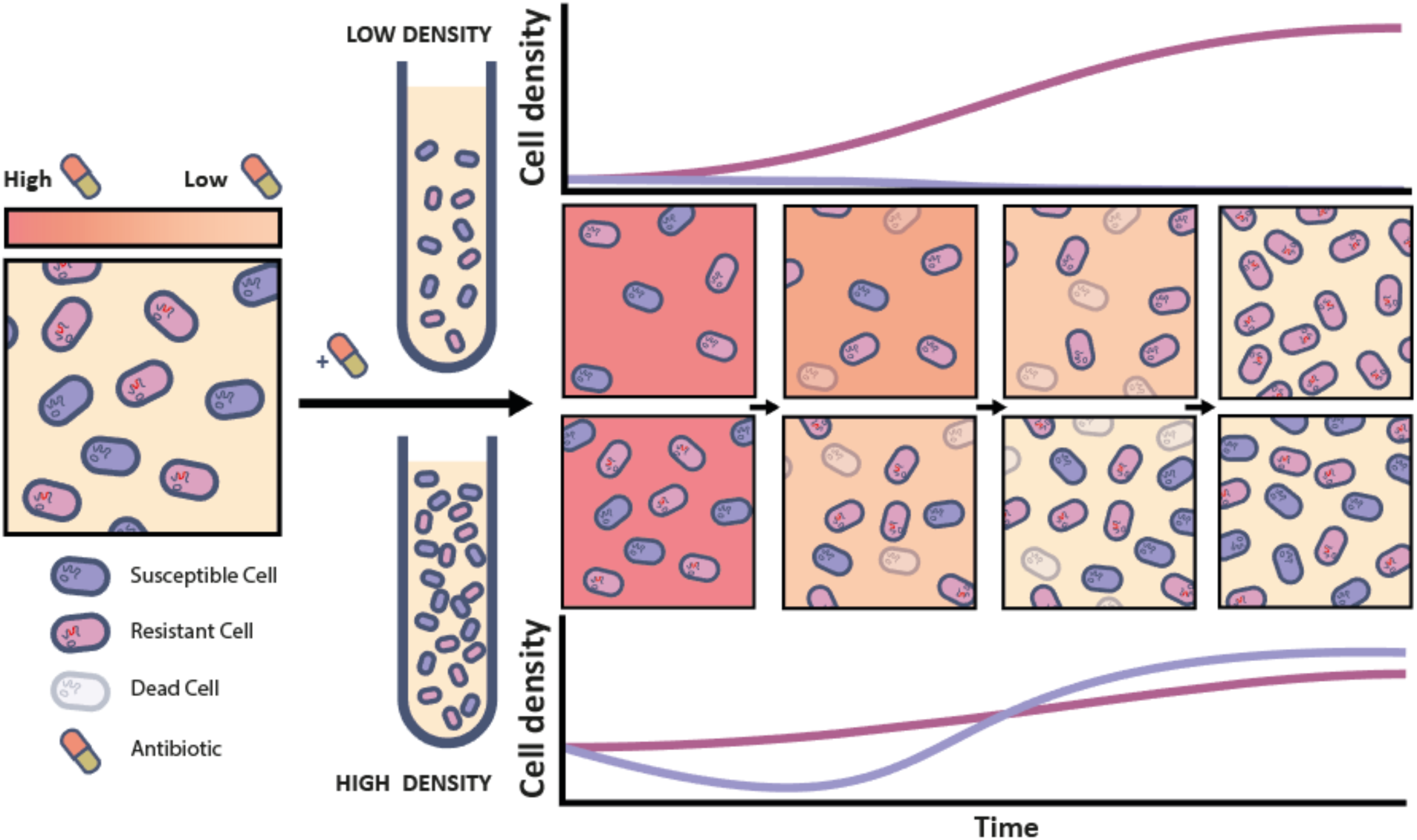
Schematic depiction of competition between resistant bacteria expressing an antibiotic-degrading enzyme (pink) and susceptible bacteria (purple) in the presence of antibiotic (background shades of red). The enzyme inactivates the antibiotic, thereby reducing the antibiotic concentration in the environment and creating conditions for susceptible bacteria to grow in the presence of resistant bacteria. The relative fitness of both strains is density-dependent, as high-density bacterial populations can degrade antibiotics more rapidly, allowing populations of susceptible bacteria to survive under otherwise lethal antibiotic concentrations.

In this study, we investigate the extent to which ecological feedback caused by antibiotic degradation can influence the spread of resistance-conferring mutations in the naturally occurring narrow-spectrum β-lactamase TEM-1. TEM-1 efficiently inactivates penicillins and first-generation cephalosporins, but just a few mutations are required to confer resistance to an extended spectrum of clinically-relevant β-lactams (21–23). To understand how resource competition and antibiotic breakdown affect the fitness of resistant relative to susceptible strains, we competed two *Escherichia coli* strains expressing distinct extended-spectrum TEM alleles which differ by two point mutations (hereafter called β-lactamase mutants) in varying concentrations of the third-generation cephalosporin cefotaxime (CTX). Specifically, we used TEM-19 (“susceptible”), which exhibits low activity against CTX, and TEM-52 (“resistant”), characterized by high activity against CTX. Both alleles are commonly observed in hospitals globally, with TEM-52 notably widespread across bacterial species (24,25) To explore the effect of eco-evolutionary feedback on the evolutionary success of mutant β-lactamases such as TEM-52, we manipulated nutrient availability to alter cell density and consequently antibiotic degradation dynamics. In addition, to quantitatively understand the experimental outcomes, we developed a population dynamic model that interrogates how physiological parameters and ecological conditions influence the relative evolutionary success of these β-lactamase mutants. Our findings highlight that ecological feedback via antibiotic degradation and resource competition critically shapes the relative fitness of the competing strains, ultimately determining the conditions under which susceptible and resistant bacteria can stably coexist.

## Results

### Population density and antibiotic concentration mediate coexistence of β-lactamase mutants

To explore the extent by which antibiotic concentration and cell density affect the fitness and possible coexistence of bacteria expressing different variants of an antibiotic-degrading enzyme, we conducted serial passage competition experiments between two isogenic strains of *E. coli* expressing different TEM β-lactamase alleles. The susceptible (TEM-19 with mutation G238S) and resistant (TEM-52 with mutations E104K, M182T, and G238S) strain have highly different MIC for cefotaxime (CTX): MIC_TEM-19_ = 1.0 μg/mL, MIC_TEM-52_ = 4,096 μg/mL (Supplementary Data 1). Thus, while both strains are able to degrade CTX more efficiently than TEM-1, they do so at highly markedly different rates (27). To distinguish them, the susceptible and resistant strain were chromosomally labelled with yellow (YFP) and blue fluorescent protein (BFP), respectively. Notably, the expression of these fluorescent proteins did not differentially affect fitness (Supplementary Figure S1A).

The maximum cell density of the cultures was modulated by supplementing glycerol as the carbon source to final concentrations of 0.2%, 0.4%, 0.9%, or 2%. Increases in glycerol concentration led to a statistically significant, although modest, increase in cell density after overnight culturing (χ² = 42.2, p < 0.0001; Supplementary Figure S1B). Furthermore, we investigated the impact of three different initial antibiotic concentrations: 12.5, 25 and 50 μg/mL CTX. We performed a total of five serial transfers using three starting ratios of TEM-52:TEM-19 (1:9, 1:1 and 9:1) and eight replicate cultures per condition and determined the fraction of resistant and susceptible bacteria after each growth cycle by counting the number of YFP and BFP cells (cells/mL) using a flow cytometer. We anticipated that if coexistence would occur, the different starting ratios would converge to a single equilibrium ratio of both strains.

Our results show that the susceptible strain (TEM-19), managed to survive at antibiotic concentrations significantly higher than its MIC in the presence of the resistant strain (TEM-52; see Figure 2A). At an antibiotic concentration of 50 μg/mL, we observed that the average final fraction of resistant cells decreased from 0.92 to 0.43 with increasing glycerol concentrations. At lower initial antibiotic concentrations of 12.5 and 25 μg/mL, varying glycerol concentrations hardly affected the final fraction of resistant cells., stabilizing at ∼0.21. However, we did observe an intriguing exception to this trend in competitions initiated with a 1:9 ratio of resistant to susceptible bacteria at glycerol concentrations of 0.2% and at 0.4% when supplemented with 50 μg/mL CTX. Here, the fraction of resistant cells surged rapidly during the initial growth cycle, overshooting the apparent equilibrium and causing the extinction of the susceptible strain.

**Figure 2.**
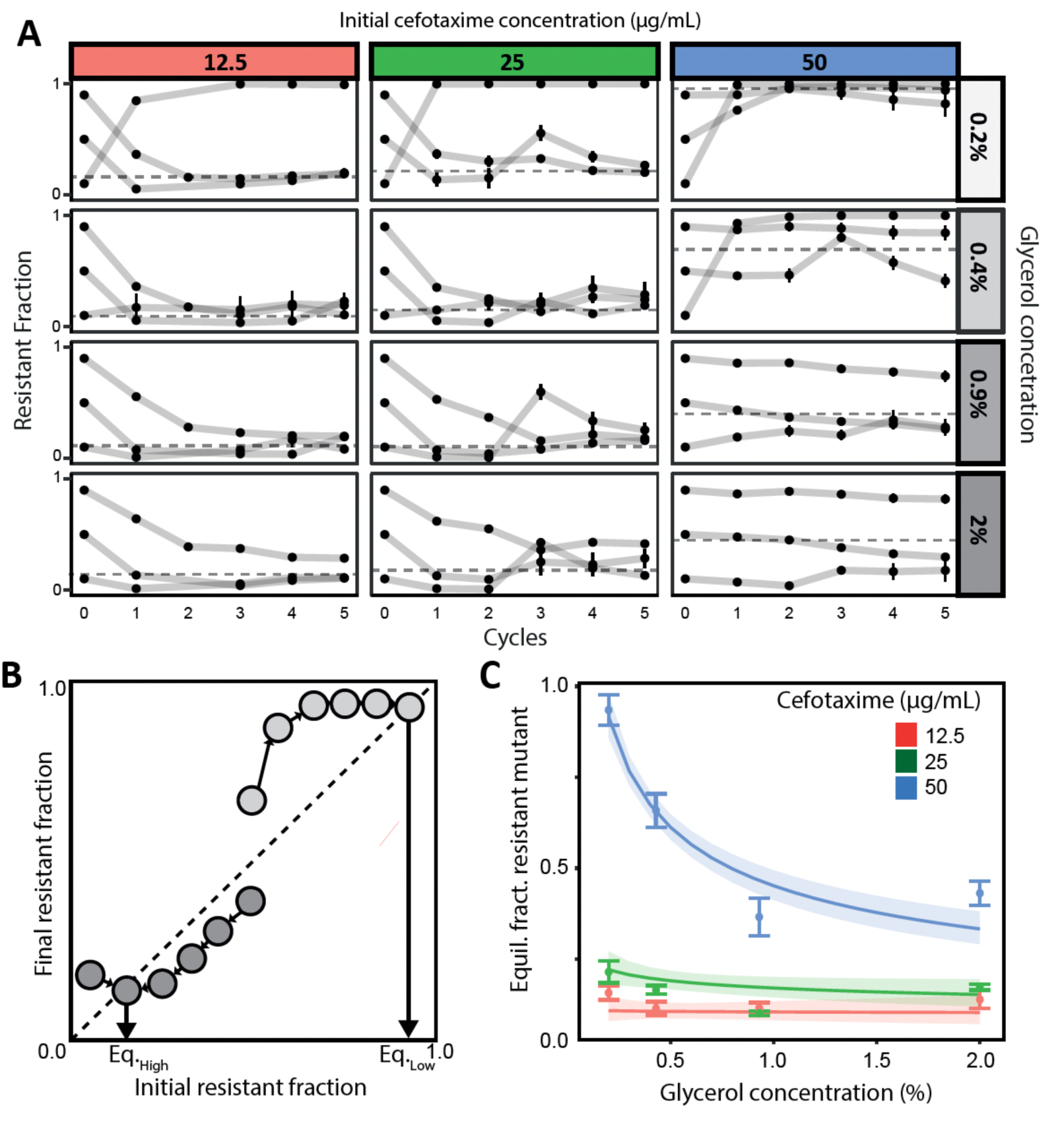
Conditions for coexistence of antibiotic-resistant and susceptible strain. **(A)** Line plots showing the fraction of the resistant strain expressing TEM-52 β-lactamase in mixed cultures together with a relatively susceptible strain expressing TEM-19 β-lactamase across five daily growth cycles for different starting ratios, cell densities (varied by varying glycerol concentrations) and CTX concentrations. The error bars represent the standard error of the mean from eight biological replicates. Note that the error bars are not always visible due to the small standard error at that point. Dotted lines show the mean of the estimated equilibrium fractions. **(B)** Schematic illustration of the procedure to infer the equilibrium frequency of the resistant strain from experimental data by plotting the resistant frequency after each growth cycle. The equilibrium is reached when the lines connecting subsequent frequencies cross the diagonal (Supplementary Section S.2). The crossing of the diagonal will be lower for high glycerol (dark grey) than for low glycerol (light gray). **(C)** Plot showing the joint impact of glycerol and CTX on the equilibrium fraction of resistant bacteria, estimated with a statistical model. The shaded areas represent the smoothed 95% confidence interval around the model estimated means (depicted by the thicker lines). Mean and standard errors of the inferred equilibria from the experiments are depicted as dots and vertical bars.

The observed convergence of strain frequencies starting from different initial fractions suggests that a dynamic equilibrium exists. In the dynamic equilibrium, strain frequencies still fluctuate across transfers, but in the same manner per transfer, leading to similar fractions at the beginning and the end of each transfer, i.e., the equilibrium ratios. Equilibrium ratios were determined with a method from Yurtsev et al. (2013) (7) for each replicate competition (Figure 2B, Supplementary Section S.2; see (7,28)). We used generalized linear models to evaluate the influence of glycerol and CTX concentration and their interaction on the equilibrium frequency of the resistant strain.

Including the interaction term significantly improved the model fit (AIC_corrected_ with interaction = −174.59; AIC_corrected_ without interaction = −168.70; Interaction effect: 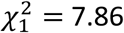, p = 0.0050). The antibiotic concentration also significantly affected the equilibrium frequency independently of glycerol (main effect antibiotic: 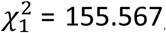, p < 0.0001), while glycerol alone did not have a significant effect (main effect glycerol: 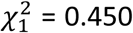, p = 0.50). Our results thus revealed that the effect of glycerol on the equilibrium frequency was modulated by the antibiotic concentration within the investigated range. In particular, glycerol, in the range tested, had little impact at low CTX levels, but significantly determined equilibrium frequencies at high antibiotic concentrations (Figure 2C).

### Physiological model predicts competitive dynamics and coexistence

We then constructed a model that could predict the competitive dynamics and coexistence of two β-lactamase-producing strains as function of antibiotic concentration and cell density, based on basic physiological parameters. We expanded a previously used model with death in stationary phase and antibiotic degradation by the more susceptible strain, two properties that we observed in our system (7). The model parameters include only the maximal growth rate (without CTX), the death rate (at CTX concentrations above MIC) and antibiotic clearance rate of each strain. The maximal growth rates (0.185±0.00326 h^-1^ for the resistant and 0.190±0.00219 h^-1^ for the susceptible strain) showed a small cost of the resistant allele, which is a necessary condition for coexistence. The death rate of the susceptible strain (1.54±0.186 h^-1^), which did not depend on the CTX concentration for concentrations used in our experiment, was higher than expected, as in previous studies it was in the order of the growth rate, while here it is an order higher (29,30). The resistant strain degraded the antibiotic faster than the susceptible strain, as expected. We found that this is caused only by increased affinity (27), because the maximal degradation rate for the susceptible strain is even higher than that of the resistant strain (6.69*10^-4^ ± 3.36*10^-4^ µg cell^-1^ h^-1^ versus 3.10 * 10^-4^ ± 9.15*10^-5^ µg cell^-1^ h^-1^ for the resistant strain). However, this maximal degradation rate of the susceptible strain is never reached due to its low affinity for CTX. We estimated these parameters in experiments (Supplementary Data 1) and used their values to parameterize our model and predict the change in the relative frequencies of the strains at each passage (see Figure 3A).

**Figure 3:**
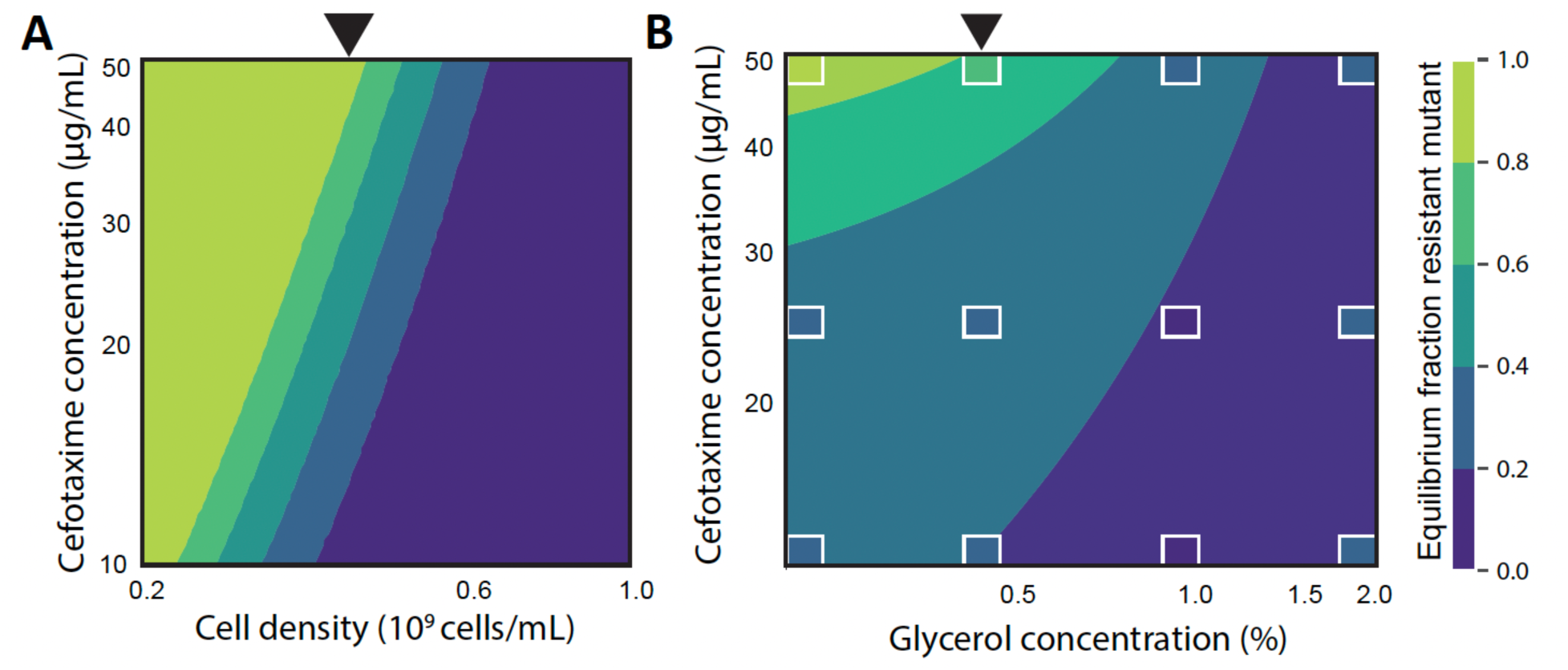
Model predictions of the equilibrium resistant frequency. **(A)** Contour plot showcasing model-projected equilibrium frequencies of the resistant strain using physiological parameter measurements from both strains, across a variety of conditions. The black arrow indicates the estimated population sizes corresponding to the experimental setup with 0.4% glycerol. **(B)** Contour plot, interpolating the statistical model represented in Figure 2C, demonstrating how the equilibrium frequencies of the resistant strain change depending on concentrations of glycerol and CTX. White boxes indicate the experimentally measured equilibrium frequencies.

When accounting for only these basic strain properties, our model predicts that a higher initial antibiotic concentration favors the resistant strain, while a higher cell density favors the susceptible strain (Figure 3A). Because the dilution rate is fixed in these simulations, increasing the cell density increases both the initial and final population size. The initial cell density affects the dynamics, because a higher initial cell density causes a faster degradation of the antibiotic, and the final cell density affects the dynamics, because a higher final cell density allows for a longer growth period once the antibiotic is sufficiently degraded. Both these density effects favor the more susceptible strain, causing the equilibrium frequency of the resistant strain to increase with antibiotic concentration and decrease with cell density (Figure 3A). Moreover, the range of CTX and glycerol concentrations that allow for coexistence largely agree with the previously described general linearized model parameterized with the experimental measurements (Figure 3A and B). It shows that for the lower tested antibiotic concentrations (12.5 and 25 μg/mL) we would need lower glycerol concentrations to see coexistence.

### Effect of cell density on coexistence in continuous culture

Whereas serial-passage experiments include large fluctuations of resource supply and population size, natural conditions often more resemble a continuous regime. Antibiotics are administrated either intermittently, with up to several doses a day, or continuously. To assess the scope for coexistence of both strains under these more clinically-relevant conditions, we modeled the intermittent regime with a continuous culture model with antibiotics added in pulses to mirror the clinical daily dosage administration of antibiotics. Blood CTX concentrations of up to 200 µg/mL are reported (31,32), depending on the weight of the person and the dosage, hence we have chosen a concentration of 100 µg/mL after administration. Similar results are obtained for continuous administration of antibiotics (see Supplementary section S.3). Under this framework, the average population size is regulated by adjusting nutrient concentrations in the inflow, while the actual population sizes fluctuate with the antibiotic pulses (Figure 4A). Consequently, culture density remains relatively constant, eliminating the initial cell density variable. However, the impact of cell density on the competitive dynamics and coexistence of both antibiotic-resistant strains remains and is qualitatively similar to our previous results. For a large range of antibiotic administration regimes, whether the susceptible wins, the resistance wins or there is coexistence depends on the cell density (Figure 4B).

**Figure 4.**
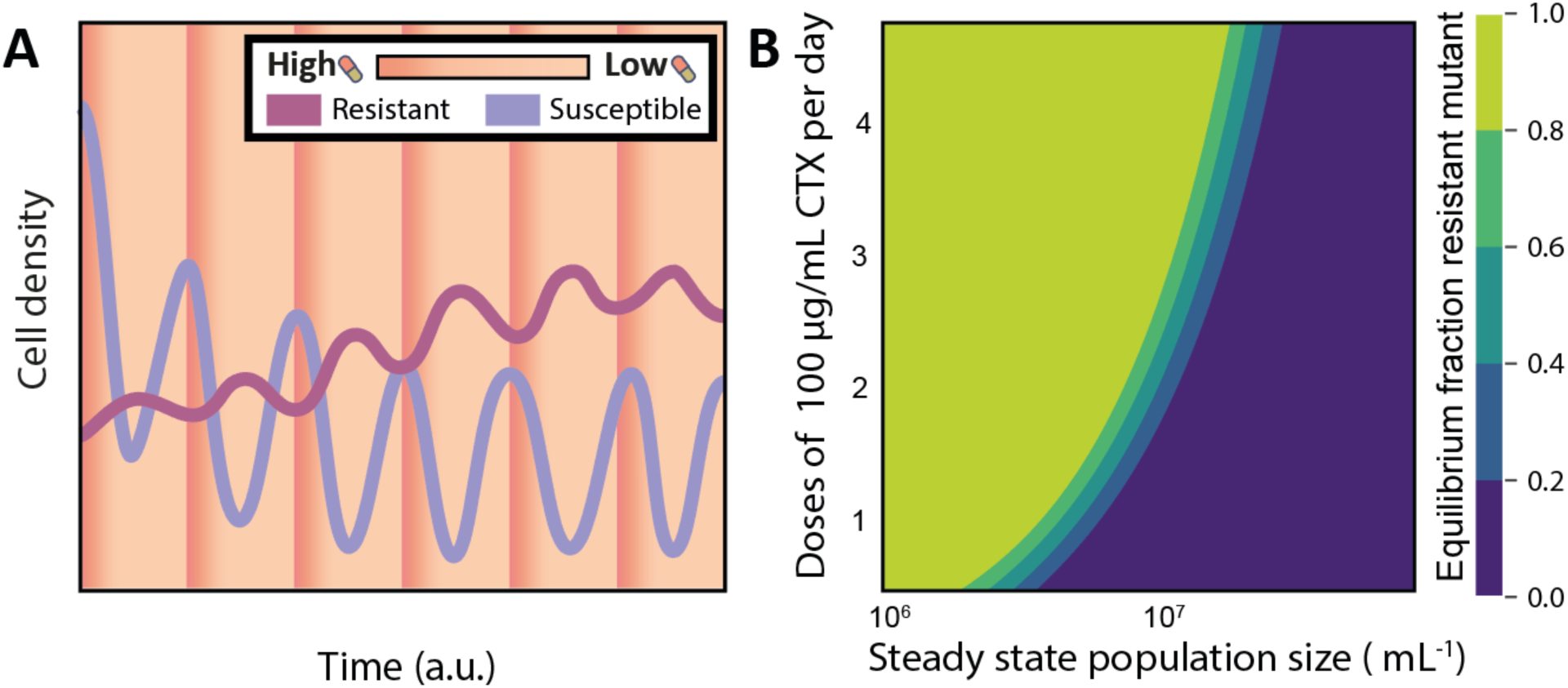
Modelling competition and coexistence under continuous nutrient input. **(A)** The model parameterized with the physiological parameters (Supplementary Data 1) and validated on the serial transfer experiments (Figure 3) is simulated in a continuous culture with antibiotic pulses (shown as shades of orange). After a pulse, the susceptible population decreases due to death and washout, which increases the nutrient availability for the resistant population, which therefore increases in size. The resistant cells degrade the antibiotic to a level at which the susceptible population recovers, which reduces the nutrient availability for the resistant population. After an initial dynamic period, the fluctuations become regular and the ratio between the strains at the start of each pulse reaches an equilibrium. **(B)** Antibiotic pulses and a constant nutrient supply leads to either a monomorphic or a mixed population of bacteria expressing different TEM β-lactamase variants. The culture density shows an effect on the equilibrium frequency of the resistant TEM variant. Cell densities and nutrients are modeled as a continuous culture and CTX is modeled as regular pulses that lead to a maximum concentration of 100 μg/mL in the culture. The growth rates of the two strains are modelled with a logistic function that depends on the CTX concentration and was experimentally obtained and a nutrient dependent term with an arbitrary but equal affinity.

## Discussion

Resistance mechanisms involving antibiotic degradation are vivid examples of ecological feedback mechanisms that are affected by their own evolution. Understanding the evolutionary role of such feedback mechanisms is essential for a comprehensive understanding of the emergence and spread of antibiotic resistance. We explored the potential for eco-evolutionary feedback via antibiotic degradation to modulate relative fitness and facilitate the coexistence of near-isogenic strains with distinct levels of antibiotic resistance. Specifically, we focused on strains carrying alleles of the antibiotic-inactivating β-lactamase enzyme TEM-1, that differed by just two mutations, resulting in large differences in their degradation efficiency of CTX and variable fitness costs in the absence of the antibiotic (20). We found that both strains showed dynamic fitness profiles that did not only depend on the antibiotic concentration, but also on the cell densities of both strains. This dynamic fitness profile ultimately dictates whether one strain dominates and becomes fixed within the population, or whether they stably coexist.

It is important to emphasize that in our study, we manipulated nutrient levels by supplementing glycerol, which only modestly affected final population density, likely due to the presence of fixed concentrations of casamino acids in the medium. Nevertheless, this variation was sufficient to alter the competitive outcome in the predicted direction. Importantly, we kept the number of generations per cycle constant, allowing us to decouple cell density from the duration of competition. This deviates from the approach taken by Yurtsev et al. (7), who demonstrated that the initial cell density, achieved by varying dilution factors, can affect the equilibrium fraction. Moreover, previous studies of coexistence via antibiotic exposure protection (33,34) typically involved coexistence between a susceptible population and a population with resistance conferred through the introduction of an antibiotic resistance gene. This approach can introduce genetic differences other than the resistance mechanism between the two populations, resulting in more pronounced fitness differences. Our results demonstrate that eco-evolutionary feedback from antibiotic degradation resulting from two single mutations can qualitatively affect the outcome of competition, with resistant mutants compromising their own selective advantage and frustrating their selective spread to high frequency. We further show that our population dynamic model, which requires no fitting parameters, can successfully predict the conditions under which coexistence arises.

Antibiotic concentrations experienced by a bacterial population are dynamic and determined by a multitude of factors, including physical environmental conditions such as temperature and pH, as well as the metabolic activities of the bacteria themselves (18). Here, we have studied the effect on the competitive success of bacteria expressing different β-lactamases, which can reduce environmental antibiotic concentrations (18,33,35). Earlier modeling work examining a small set of TEM-1 β-lactamase mutants found that the evolution of CTX resistance via mutations enhancing CTX degradation is dependent on the administered antibiotic concentrations (36). In another study, the coexistence of susceptible and plasmid-mediated ampicillin resistant populations has been shown to lead to the evolution of the susceptible strain towards greater antibiotic tolerance (less death with antibiotic exposure) without significant increases in resistance (no increase in growth with antibiotic exposure), unlike monocultures where resistance would increase (8). Our study demonstrates that antibiotic exposure protection also leads to stable coexistence among closely-related TEM-1 alleles, indicating the importance of eco-evolutionary feedback for the evolution of β-lactamase alleles with enhanced activity against the β-lactam used for selection. Our findings demonstrate that predictions of evolutionary trajectories of antibiotic resistance relying on traditional resistance measurements, such as MIC (6), do not accurately capture the complex dynamics governing resistance evolution under all conditions (37). Instead, population dynamic models capturing effects from antibiotic degradation on relative fitness become imperative to account for unavoidable eco-evolutionary feedback processes.

Our experiments revealed that in some cases, there was an unexpectedly marked increase in the resistant fraction, at times even leading to the extinction of the susceptible population, especially when the resistant fraction is initially low. This phenomenon can be attributed to accidental low numbers of transferred resistant cells, which keep the antibiotic concentration at elevated levels for an extended period. As a result, the susceptible population experiences a rapid and substantial decline that cannot be compensated for by subsequent growth. This effect of overshooting the equilibrium, which we observed in our experiments, has also been reported in a previous study (7), and is replicated in our model, underscoring its validity.

We modified a population dynamic model based only on very basic physiological parameters. Our results with this model show that we can largely understand the quantitative outcomes of the experiments based on the properties of the individual strains, which we have validated with experimental data. Our model’s ability to predict coexistence without the need for parameter estimation during model fitting (in contrast to previous similar approaches (e.g. (7)), indicates that our model successfully captured the most relevant factors for predicting the effect of ecological feedback in various scenarios (see Supplementary Figures S5-S7). Importantly, our model allowed us to make predictions beyond the setting of the experiments to more clinically relevant conditions. Using a continuous culture with pulsed antibiotic administration, we demonstrated that the observed patterns hold true in more realistic scenarios.

The density effect on coexistence that we describe in this paper is expected to be relevant also for other microbial interactions. Most closely, resistance conferred by other antibiotic-degrading enzymes, including other β-lactamases and acetyltransferases capable of inactivating aminoglycosides or chloramphenicol (38), is likely to involve similar feedback on bacterial fitness as discussed for the TEM mutants. However, population density also affects metabolic interactions: cross-feeding is only beneficial at intermediate densities and production of compounds that are self-inhibitory can lead to coexistence depending on the density (39,40). Therefore, much is to be learned about population density effects from feedback mechanisms on the coexistence and evolution of microorganisms in microbial communities.

While our model successfully captures the feedback effect of antibiotic degradation on the relative fitness of a β-lactamase mutant, it has limitations compared to natural populations and conditions. One of the limitations is that our study is limited to mutations enhancing the activity of the β-lactamase and does not consider compensatory mutations that can reduce the costs of resistance, which may reduce the potential for cross-protection to frustrate the evolution of antibiotic resistance. Nor do we consider other, more private, mechanisms of resistance, such as target modification or decreased intake or increased efflux of antibiotics, which will potentially come with different forms of eco-evolutionary feedback (41,42). Expanding our model to include spatially-structured conditions, such as those found in biofilms, is also important. These environments are more representative of natural conditions and may significantly influence the dynamics we have observed due to strong local feedback effects (11,34,43). Nevertheless, our study provides a mechanistic framework for predicting how ecological feedback may influence the evolution of resistance via antibiotic degradation – a resistance mechanism allowing for strong feedback. Integrating such dynamics into experimental and clinical models will be essential for developing effective, evolution-informed strategies to contain antibiotic resistance.

## Methods

### Strains, plasmids and medium

The *Escherichia coli* strain used throughout this study was BW27783 (CGSC#12119), which lacks arabinose metabolizing genes and expresses *araE* from an arabinose independent promoter allowing stringent control of the arabinose-inducible promoter, pBAD (44). Plasmid pBAD332T was used as the expression vector to express TEM-1 and its alleles, TEM-19 (G238S) and TEM-52 (E104K-M182T-G238S)(45). To differentiate between the two different alleles during the coexistence experiment, BW27783 was labelled with a yellow (YFP) or blue (BFP) fluorescent marker by inserting cat-J23101-SYFP2 or cat-J23101-mTagBFP2 into chromosomal *galK* using the Quick and Easy *E. coli* Gene Deletion Kit (Gene Bridges, Heidelberg, Germany). We evaluated the fitness effect of the fluorescent proteins through pairwise competition assays. Chromosomally labeled YFP and BFP cultures were mixed and diluted at a 1:1,000 ratio in fresh M9 minimal medium(n=24). Cell frequencies were measured at t=0h and t=24h using flow cytometry. Relative fitness was calculated as the ratio of Malthusian parameters (46).

TEM alleles were amplified from previously described plasmid constructs (47), ligated into pBAD322T using T4 DNA ligase (New England Biolabs), and then electroporated into *E. coli* DH5α. After verifying successful cloning via sequencing, isolated plasmid was transformed into BW27783, BW27783-YFP and BW27783-BFP. All experiments were performed in M9 minimal media (supplemented with 0.2% Casamino acids, 1µg/mL thiamine, and 10 µg/ml tetracycline to prevent plasmid loss). Glycerol was added to the medium as carbon source to a final concentration of 0.4% unless otherwise stated. L-arabinose (0.000625% or otherwise stated) was used to express the TEM-alleles.

### Growth rate, death rate and minimum inhibitory concentration

To determine the growth rate of BW27783 expressing TEM alleles TEM-19 or TEM-52, 200 µL M9 medium supplemented with cefotaxime and L-arabinose in a 96-well microtiter plate was inoculated at a 1:200 dilution from overnight culture in M9. The samples were subsequently covered with 50 µL mineral oil to prevent evaporation and condensation of the lid. The 96-well plate was incubated for 24h at 37 °C in a Victor3 microtiter plate reader (PerkinElmer, Waltham, MA, USA), and absorbance was measured every three minutes. The plate was shaken (double orbital) for 30 seconds before each measurement. Growth rates represent the maximum linear slope of the natural logarithm of absorbance versus time in a 2 h window and were determined using a custom python script.

The death rate of BW27783 expressing TEM-19 was estimated by performing a time-kill experiment. Bacteria from an overnight culture grown in M9 medium supplemented with L-arabinose were exposed to 25 µg/ml cefotaxime. After 0, 2 and 6 hours of incubation at 37°C, the number of colony forming units per ml were determined by plating on LB agar plates and counting the number of colonies formed after overnight incubation. The death rate was calculated as the slope of the regression line describing the logarithm of the bacterial density as a function of time.

We determined the minimum inhibitory concentration of cefotaxime of both TEM mutants using the liquid broth microdilution method (48). From overnight culture in M9 medium, approximately 200,000 cells were transferred to 200µl of M9 medium supplemented with cefotaxime and L-arabinose. Plates were incubated for 24 h at 37 °C with agitation. The minimum inhibitory concentration was visually assessed and defined as the lowest cefotaxime concentration where no growth could be observed.

Growth rate and minimum inhibitory concertation were determined for BW27783 carrying pBAD322T-TEM-19 and BW27783 with pBAD322T-TEM-52. Death rate was only determined of BW27783 expressing TEM-19.

### Estimating the in vivo cefotaxime degradation rate

To qualitatively asses the *in vivo* degradation rate of both β-lactamase alleles, we developed a bioassay based on the agar diffusion assay (49). In this assay, we estimated the remaining cefotaxime concentration in medium inoculated with BW27783 carrying either pBAD322T-G238S or pBAD322T-E104K-M182T-G238S after defined time intervals. Each strain was initially cultured overnight in 5ml M9 medium supplemented with 10 μg/ml tetracycline and 0.000625% arabinose at 37°C with continuous shaking at 250 rpm. After 16 hours of incubation, the OD600 of these samples were measured in a spectrophotometer and the cells were pelleted and resuspended in 5ml M9 medium with cefotaxime but without glycerol. The initial concentration cefotaxime for the TEM-19 mutant was 32 μg/ml and for TEM-52 4096 μg/ml. These samples were incubated at 37°C and, after 15-, 30- and 45 minutes, cell-free supernatants (CFS) were prepared and placed on ice. The CFS was prepared by pelleting down 1.4 ml culture and filter sterilizing the supernatants using a 0.22 μm filter. From the CFS of each mutant, an eight times two-fold dilutions series was made in M9 medium. At each time point, the number of colony-forming units (CFUs) was determined by collecting 1 ml culture of each mutant after 0, 15, 30 and 45 min by plating a 10,000-fold dilution on LB-agar plates and counting the number of colonies following incubation overnight.

In order to assess the remaining concentration of cefotaxime at different time points, a control dilution series with known cefotaxime concentrations was also prepared. LB-agar plates were streaked with 100 μl of overnight culture of the cefotaxime susceptible WT BW27783. Four holes were punctured in the agar arranged in equidistant positions with a pipette tip (P100). Each hole was filled with 100 μl from the control dilution series (in triplicate) or 100 μl from the CFS-dilutions series of each mutant (in triplicate) and incubated at 37°C for one day. As the susceptible WT strain cannot grow at high CTX concentrations, a zone of inhibition (ZOI) was visible on the plates of which the diameter was measured using a ruler. A calibration curve was prepared by plotting the known CTX concentrations of the control dilution series and its measured ZOI. The resulting equation was subsequently used to predict the amount of remaining CTX based on the ZOI. For each time point, the predicted concentration was divided by the average CFUs between the current time point and the previous one. The result of this was plotted against time, resulting in a linear relationship. The slope of this line represents the estimated degradation rate (ug cell^-1^ h^-1^)

### Coexistence experiment

To investigate the impact of population density and antibiotics on two strains carrying either TEM-19 or TEM-52, which were chromosomally labeled with yellow or blue fluorescent protein, we conducted a serial passage competition experiment. The induced overnight cultures were counted using a flow cytometer and mixed to three starting ratios (1:9, 1:1, and 9:1) in eight biological replicates. Then, these mixed cultures were transferred to 96-well plates containing M9 medium supplemented with glycerol (0.2, 0.43, 0.93, or 2%) and cefotaxime (50, 25, or 12.5 μg/ml) to a final volume of 200 μl. The starting population size of each condition was 200 times less than the maximum cell number in the respective conditions. To ensure that the culture was fully saturated, we incubated it for 48 hours at 37°C. After 48 hours of incubation with shaking, each culture was diluted 20-fold and transferred to a new 96-well plate with freshly prepared M9 medium supplemented with glycerol and cefotaxime at the appropriate concentrations, and this serial transfer was repeated five times. Following each growth cycle, each culture was diluted 100-fold in a new plate containing cold M9-medium before being counted using a MACSQuant10 flow cytometer (Miltenyi biotec, Bergisch Gladbach, Germany). We conducted flow cytometry analysis with a moderate flow rate using SSC 1.4 as a trigger, and we counted a total of 20,000 events for each sample. For obtaining the equilibrium ratio from the experimental data, see Supplementary Section S.2.

Data analysis was conducted using R statistical software (v3.6.1) accessed through RStudio (Build 576). To assess the relationship between the equilibrium fraction and antibiotic concentration and density, a generalized linear model was applied using the ‘glm’ function (base R package). Model selection was based on the second-order Akaike information criterion using ‘AICc’ implemented in the ‘MuMIn’ package.

### Model description

We follow the modeling approach from (7), extending their Model 7 with two additions. We observe death in the lag phase and our susceptible strain also degrades the antibiotic. Therefore, we have included a death term in the lag phase and degradation by the susceptible strain. Including either of these features makes it impossible to directly calculate the equilibrium fractions, therefore we have resorted to numerical methods to obtain the equilibrium fractions (see supplementary Jupyter notebook).

Our model contains the following set of equations:

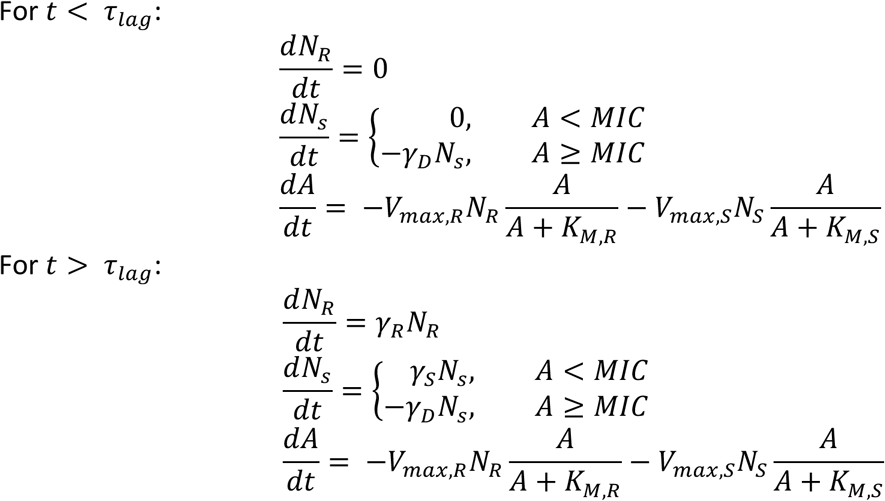

With *N_R_* the population size of the more resistant strain TEM-52, *N_S_* of the more susceptible strain TEM-19 and *A* the antibiotic concentration. For the parameters see Supplementary Table S1. See the supplementary information for the sensitivity analysis and the continuous culture model.

## Acknowledgments

We thank Marco Lezzerini for experimental assistance. This study was supported by grants from the Human Frontiers in Science Program (RGP0010/2015) and the Netherlands organization for scientific research (OCENW.XS.058) to JAGMdV and an EMBO fellowship (ALTF 273-2017) to PR.

## Declaration of interests

The authors declare no competing interests.

## Appendices

**Supplementary Information:** Supporting information including Supplementary Section S.1 (Competitive fitness and effect of glycerol concentration), S.2 (Equilibrium Ratio from Experimental Data), S.3 (Serial Transfer Model), and S.4 (Continuous Culture Model), Supplementary Tables S1, S2, and S3, as well as Supplementary Figures S1 through S7.

**Supplementary Data 1**

**Supplementary Code:** https://github.com/WortelLab/tem1coex/

## Notes

https://github.com/WortelLab/tem1coex/

### Competing Interest Statement

The authors have declared no competing interest.

### Summary of Updates

Updated text and figures, supplementary information and Code. No change to the underlying data but the representation and additional supplementary figure 1B and additional simulation in the supplementary information.

