## Supplemental Information for "Density-dependent feedback limits the spread of a beta-lactamase mutant: experimental observations and population dynamic model"

### S.1 Competitive fitness and effect of glycerol concentration

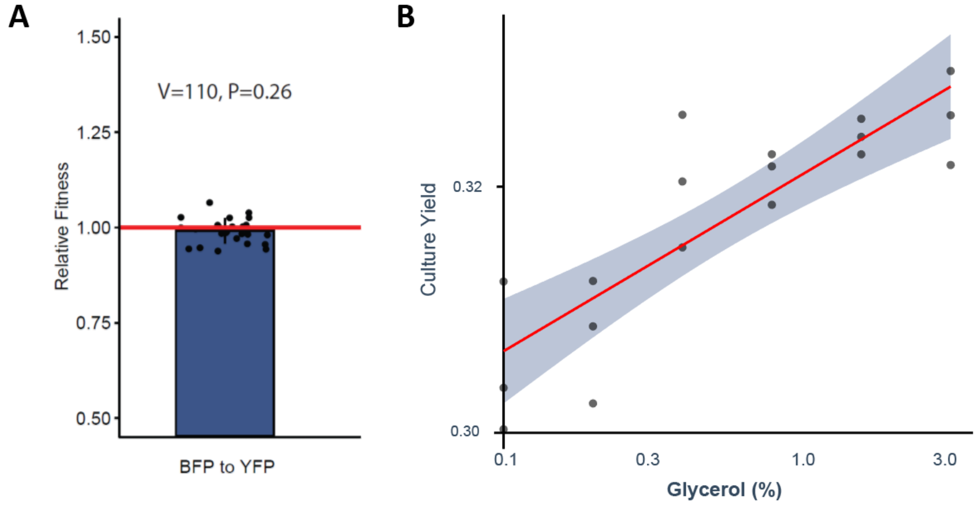

***Figure S1. A*** *Relative competitive fitness of E. coli strain BW27783 expressing fluorophore BFP to BW27783 expressing YFP. The statistical insignificance was assessed using the one-sample Wilcoxon non-parametric test (n=24).* ***B*** *Relationship between glycerol concentration and culture yield. Yield was approximated using the Gompertz model parameter K, obtained by fitting a Gompertz model to the growth curve. A Generalized Linear Model was applied to assess the effect of glycerol concentration on K, revealing a significant effect ($\chi 2=42.228$, $p<0.0001$). The solid line represents the model fit, and the shaded ribbon indicates the 95% confidence interval.*

### S.2 Equilibrium ratio from experimental data

We monitored the resistant and susceptible cells over several transfers, but at the final transfer, the equilibrium ratio might not have been reached. To obtain the equilibrium ratio from the experimental data, we have taken the approach to plot the initial versus the final ratio of a transfer. This approach is also known plotting the difference equation or 'cobweb diagrams' and used in (Yurtsev et al. 2013; Nev et al. 2020). In the equilibrium, the initial ratio is per definition equal to final ratio. Since the experiment adds noise, the experimental data will not lie exactly on the diagonal, therefore we use this approach: we sort the pairs of initial ratio and final ratio by the initial ratio and then check if any subsequent pairs cross the diagonal. Due to experimental noise, there could also be several crossings of the diagonal, in which case we take the average of those as our best estimate of the equilibrium ratio.

In some cases, the susceptible strain was lost at the first transfer, because its initial frequency was low, and the antibiotics remained high for a long time. This caused such a severe decrease in growth of the susceptible strain that the strain was lost during the bottleneck imposed at the transfer. Those cases were left out for the calculation of the equilibrium frequency, because this was the result of stochastic effects that lead to the unstable equilibrium of only resistant cells, and not to the other stable equilibrium that was inferred from the other initial ratios (these cases were a glycerol concentration of 0.2% and initial frequency of the resistant strain of 0.1 for all CTX concentrations and also glycerol = 0.43% initial frequency resistant of 0.1 and CTX=50).

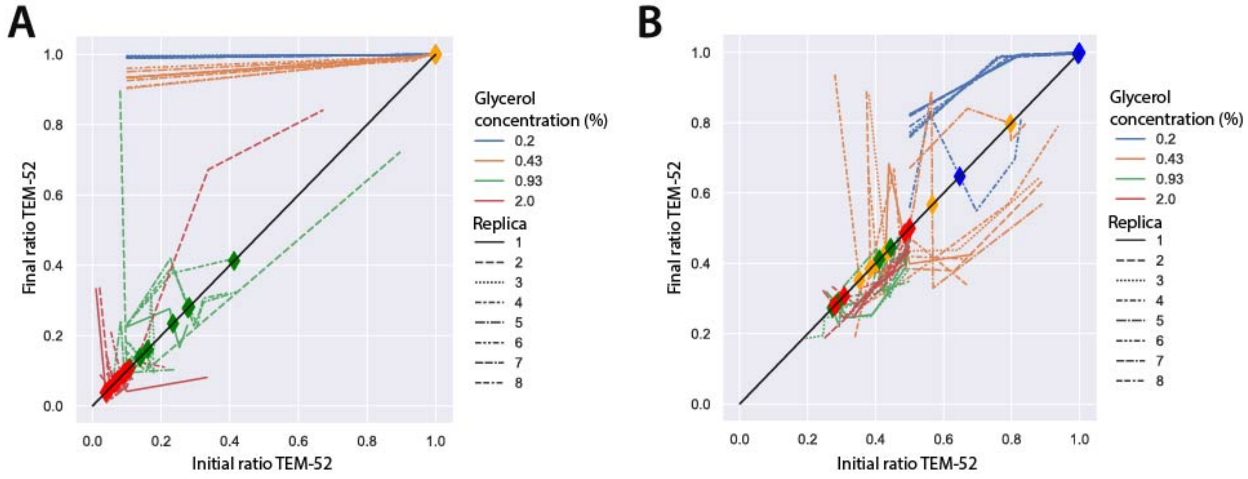

***Figure S2.*** *Example of the outcome of the calculation of the crossing of the diagonal. Lines connect pairs of (initial ratio, final ratio) for one experiment. Diamonds show the calculated equilibrium frequency (crossing of the diagonal). Note that the order of the pairs is by initial ratio, not necessarily by cycle number. For the cycles, a subsequent cycle necessarily has the initial ratio equal to the final ratio of the previous cycle. For example, the green line in **A** starting at around (0.1, 0.9) at a certain cycle, has the next cycle at the point around (0.9, 0.72). **A** CTX=50 and initial ratio = 0.1. For higher densities (higher glycerol concentrations) we get a coexistence with more of the susceptible strain. We dropped the data for the two lowest concentrations of glycerol for the statistical model fit, because the resistance strain was quickly lost due to its low initial frequency and the bottleneck. **B** Similar graph for a starting ratio of 0.5 of the resistant strain. In this case the strains were not lost after the first transfer.*

### S.3 Serial transfer model

#### S.2.1 Model parameters

The parameters for the serial transfer model are inferred from our measurements, except for the affinity of the degradation of CTX, which is taken from literature. For the lag time and the yield, an estimate is made on the basis of our data. All parameters are shown in Table S1.

| Parameter | Description | Value (± St dev) | Unit | Obtained from |
|---|---|---|---|---|
| $\gamma_D$ | Death rate TEM-19 | 1.54±0.186 | $h^{-1}$ | Bottom value from fit of growth rates at different CTX values and death rate assay |
| $\gamma_R$ | Growth rate TEM-52 | 0.185±0.00326 | $h^{-1}$ | Average of measurements at 0 CTX |
| $\gamma_S$ | Growth rate TEM-19 | 0.190±0.00219 | $h^{-1}$ | Average of measurements at 0 CTX |
| MIC | Minimal inhibitory concentration TEM-19 | 1 | μg $ml^{-1}$ | Minimum inhibitory concentration |
| $V_{max,R}$ | Maximal hydrolysis rate TEM-52 | $3.10 * 10^{-4} \pm 9.15*10^{-5}$ | μg $cell^{-1}$ $h^{-1}$ | Degradation rate assay |
| $K_{M,R}$ | Affinity hydrolysis TEM-52 | 69 | μg $ml^{-1}$ | (Knies, Cai, and Weinreich 2017) |
| $V_{max,S}$ | Maximal hydrolysis rate TEM-19 | $6.69*10^{-4} \pm 3.36*10^{-4}$ | μg $cell^{-1}$ $h^{-1}$ | Degradation rate assay |
| $K_{M,S}$ | Affinity hydrolysis TEM-19 | 319 | μg $ml^{-1}$ | (Knies, Cai, and Weinreich 2017) |
| $\tau_{lag}$ | Time before onset of growth | 8.5 | h | Estimate from our data |
| Yield | Cell yield on glycerol | $2 * 10^9$ | Cells (w/v glycerol) $^{-1}$ | Estimated from our data |

***Table S1.*** *Parameters used in the computational model for the simulation of the serial transfers.*

#### S.2.2 Sensitivity analysis

The measurement errors were used to define appropriate distributions for the sampling of the parameters (see Table S2). 1000 parameters sets were drawn and checked for distributions and correlations (see Jupyter notebook). Then time simulations were done for all of these sets, and the 95% confidence intervals were determined (see Figure S3).

| Parameter | Value (± Standard deviation) | Unit | Distribution |
|---|---|---|---|
| $\gamma_D$ | 1.54 ± 0.186 | $h^{-1}$ | Normal distribution |
| $\gamma_R$ | 0.185 ± 0..00326 | $h^{-1}$ | Normal distribution |
| $\gamma_S$ | 0.190 ± 0.00219 | $h^{-1}$ | Normal distribution |
| MIC | 1 | μg $ml^{-1}$ | Lognormal distribution with stdev = 0.5 |
| $V_{max,R}$ | $3.20 * 10^{-4} \pm 9.46 * 10^{-5}$ | μg $cell^{-1}$ $h^{-1}$ | Normal distribution |
| $K_{M,R}$ | 69 ± 7 | μg $ml^{-1}$ | Lognormal distribution |
| $V_{max,S}$ | $6.69 * 10^{-4} \pm 3.36 * 10^{-4}$ | μg $cell^{-1}$ $h^{-1}$ | Normal distribution |

| | | | |
|---|---|---|---|
| $K_{M,S}$ | 319 ± 182 | μg ml$^{-1}$ | Log-normal distibution |
| $\tau_{lag}$ | 8.5 | h | Normal with stdev of 1 |
| Yield | 2 * $10^9$ | Cells (w/v glycerol) $^{-1}$ | Normal with stdev of 1* 10^9 |

***Table S2.*** *Parameter distributions used for the sensitivity analysis*

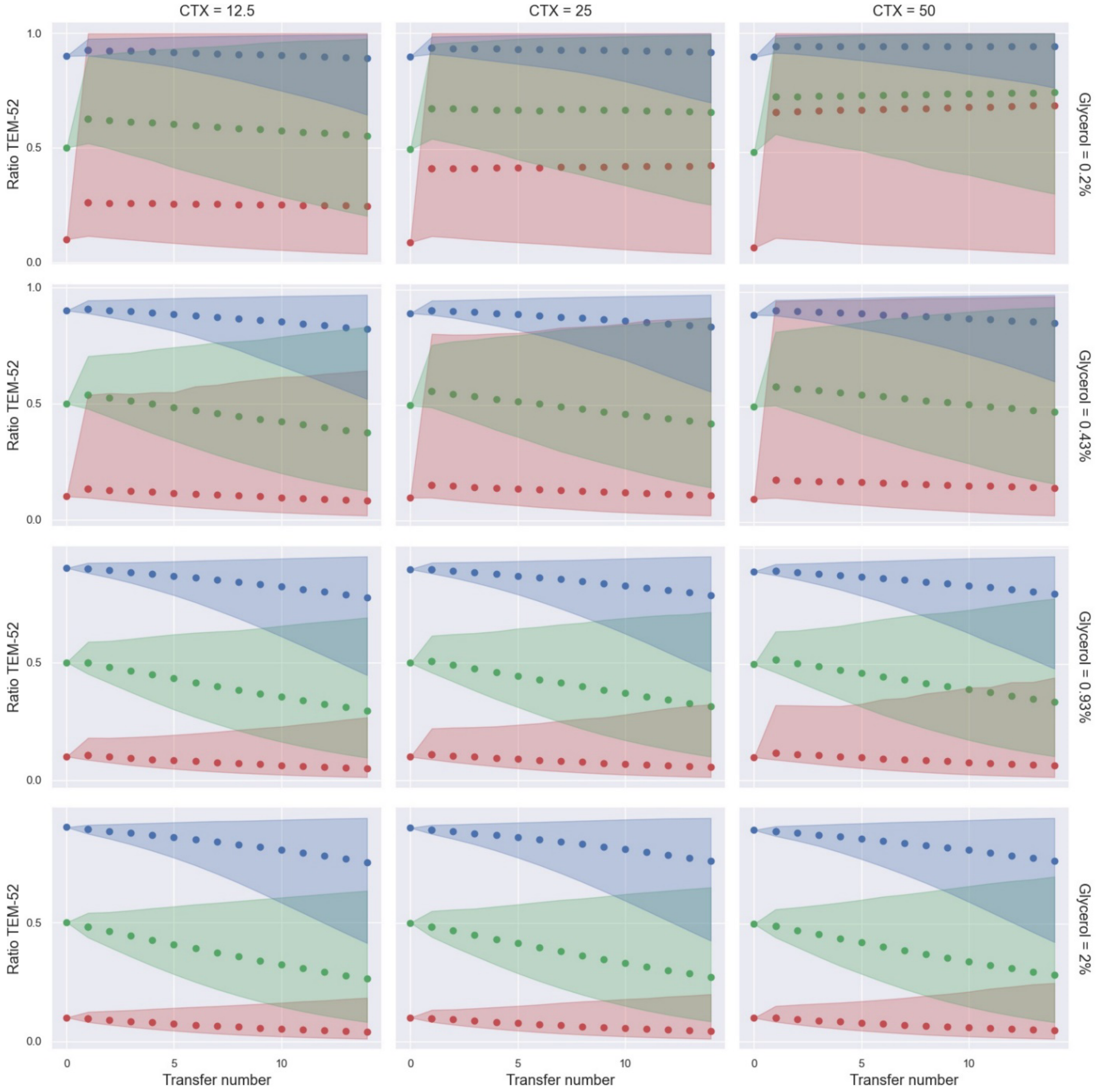

***Figure S3.*** *Simulations of the serial transfers at the different glycerol and Cefotaxime concentrations used in the experiment. Dots show the frequency of the resistant strain at the time of the transfer, and the cones show the 95% confidence intervals.*

### S.4 Continuous culture model

#### S4.1 Description of the model

The differential equations for the continuous culture are:

$$\frac{dg}{dt} = f_g * d - \frac{1}{y}\left(N_R\ \mu_R(\mathrm{A})\frac{g}{k+g} + \ N_S\ \mu_S(\mathrm{A})\frac{g}{k+g}\right) - g * d$$

$$\frac{dA}{dt} = f_A * d - V_{max,R} N_R \frac{A}{A + K_{M,R}} - V_{max,S} N_S \frac{A}{A + K_{M,S}} - A * d$$

$$\frac{dN_R}{dt} = \ N_R\ \mu_R\ \frac{g}{k+g} - d * N_R$$

$$\frac{dN_S}{dt} = \ N_S\ \mu_S\ \frac{g}{k+g} - d * N_S$$

$g$ is the glycerol concentration in the culture, $f_g$ the glycerol concentration in the feed, $f_A$ the antibiotic concertation in the feed and $d$ the dilution rate.

#### S4.2 Parameters

The additional parameters used for the continuous model are listed in Table S3. For the fit of the growth rate at different CTX concentrations we used a 4-parameter logistic function:

$$B + \frac{T - B}{1 + \left(\frac{a}{ec50}\right)^h} \tag{1}$$

Where B is the death rate, T the maximum growth rate, ec50 the inflection point and h the hill coefficient (see Figure S4 for the fit).

| parameter | meaning | value | unit | experiment |
|---|---|---|---|---|
| $T_R$ | Top of CTX dependent growth curve (TEM-52) | 0.146 | $h^{-1}$ | Top value from fit of growth rates at different CTX values and death rate assay |
| $B_R$ | Bottom of CTX dependent growth curve (TEM-52) | -96.5 | $h^{-1}$ | Bottom value from fit of growth rates at different CTX values and death rate assay |
| $h_R$ | Hill coefficient (TEM-52) | 3.38 | | Hill value from fit of growth rates at different CTX values and death rate assay |
| $ec50_R$ | Inflection point (TEM-52) | 27716 | µg $ml^{-1}$ | Ec50 value from fit of growth rates at different CTX values and death rate assay |
| $T_S$ | Top of CTX dependent growth curve (TEM-19) | 0.183 | $h^{-1}$ | Top value from fit of growth rates at different CTX values and death rate assay |
| $B_S$ | Bottom of CTX dependent growth curve (TEM-19) | -1.57 | $h^{-1}$ | Bottom value from fit of growth rates at different CTX values and death rate assay |
| $h_S$ | Hill coefficient (TEM-19) | 3.42 | | Hill value from fit of growth rates at different CTX values and death rate assay |
| $ec50_S$ | Inflection point (TEM-19) | 1.87 | µg $ml^{-1}$ | Ec50 value from fit of growth rates at different CTX values and death rate assay |
| $k$ | Substrate affinity | $1.8 * 10^{-4}$ | % Glycerol | Converted from (Truniger and Boos 1993) |

***Table S3.*** *Additional parameters for continuous culture*

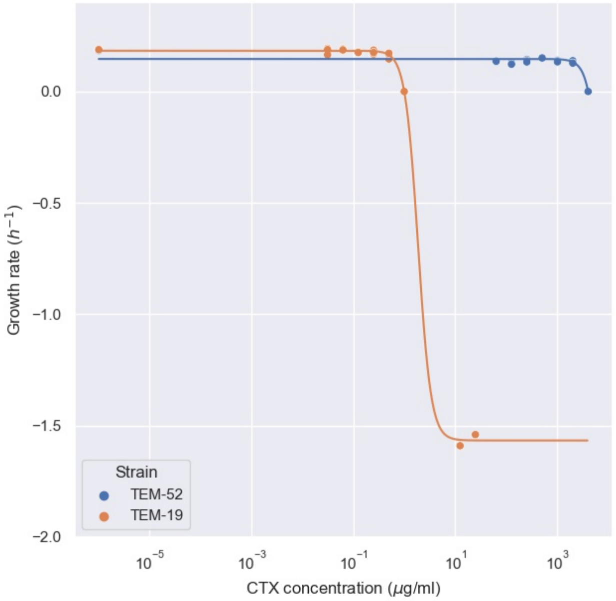

***Figure S4.*** *Fit of equation (1) to the growth data of the two strains.*

#### S4.3 Calculation of the coexistence area

To obtain the feed concentrations and dilution rate for which the two strains coexist in steady state, we use the following procedure:

Strains can only coexist when their growth rates are equal. In our set this is the case when $\mu_R = \mu_S$ , which happens when $A_{coex} = 0.61$ (see supplementary Jupyter notebook). This means if the antibiotic in the feed is lower, the susceptible strain will always win. When the concentration is higher, coexistence requires that the antibiotic is degraded to a lower level when only the Resistant strain is present, and to a higher level when only the Susceptible strain is present. This depends on the dilution rate and the glycerol and antibiotic in the feed.

For a fixed dilution rate, we can calculate the antibiotic and glycerol concentrations in the feed on the boundaries of the coexistence area. First, we find the boundary between the area where the resistant strain wins and the coexistence. From setting the differential equation of $A$ to 0 we obtain the minimal cell density of the resistant strain to achieve $A_{coex} = 0.61$, depending on $f_A$:

$$N_R = \frac{d\,(f_A - A_{coex})(A_{coex} + K_{M,R})}{A_{coex}\,V_{max,R}}$$

From the steady state of the differential equation of the resistant strain ($dN_R/dt$) we know that the growth has to be equal to the dilution rate, and therefore:

$$g = \frac{d\,k}{\mu_R(A_{coex}) - d}$$

Finally, we use the differential equation for $g$ to calculate the boundary in terms of $f_A$ and $f_g$:

$$f_{g(R-coex)} = \left(\frac{1}{y}\,\frac{(f_A - A_{coex})\left(A_{coex} + K_{M,R}\right)}{A_{coex}\,V_{max,R}}\,\frac{\mu_R(A_{coex})}{k + \frac{d\,k}{\mu_R(A_{coex}) - d}} + 1\right)\frac{d\,k}{\mu_R(A_{coex}) - d}$$

Below this line, only the resistant strain survives, because the antibiotic is too high when only the resistant is present for the susceptible strain to reach a high enough growth rate to equal the dilution rate.

Next, in a similar fashion, we calculate the boundary between the coexistence area and where only the susceptible survives. The susceptible dominates when it can alone degrade the antibiotic below $A_{coex}$. We again calculate the biomass necessary for that from the differential equation of $A$:

$$N_S = \frac{d\,(f_A - A_{coex})(A_{coex} + K_{M,S})}{A_{coex}\,V_{max,S}}$$

We can use the same calculation the glycerol concentration and then use the differential equation for $g$ to calculate the boundary in terms of $f_A$ and $f_g$:

$$f_{g(coex-S)} = \left(\frac{1}{y}\,\frac{(f_A - A_{coex})\left(A_{coex} + K_{M,S}\right)}{A_{coex}\,V_{max,S}}\,\frac{\mu_S(A_{coex})}{k + \frac{d\,k}{\mu_S(A_{coex}) - d}} + 1\right)\frac{d\,k}{\mu_S(A_{coex}) - d}$$

In the coexistence area we can calculate the equilibrium frequency of the resistant strain. In coexistence, $A = A_{coex}$ and $g = \frac{d\,k}{\mu_R(A_{coex}) - d} = \frac{d\,k}{\mu_S(A_{coex}) - d}$. We define $\mu_{coex} = \mu_S(A_{coex}) = \mu_R(A_{coex})$. Then we can calculate $N_{tot} = N_R + N_S$ from the steady state of differential equation of the substrate:

$$N_{tot,SS} = \frac{k + g}{g\,\mu_{coex}}\left(f_g - g\right)d\,y$$

Next, we can use the steady state of the differential equation for the antibiotics to calculate the steady state density of the resistant strain:

$$N_{R,SS} = \frac{(f_A - A_{coex})\,d\;V_{max,S}\,N_{tot}\,\frac{A_{coex}}{A_{coex} + K_{M,S}}}{V_{max,R}\,\frac{A_{coex}}{A_{coex} + K_{M,R}} - V_{max,S}\,\frac{A_{coex}}{A_{coex} + K_{M,S}}}$$

Using $N_{R,SS}$ and $N_{tot,SS}$ we calculate the steady state ration of the resistant strain for different substrate and antibiotic feed concentrations.

The predicted coexistence areas depend on the dilution rate (see Figure S5).

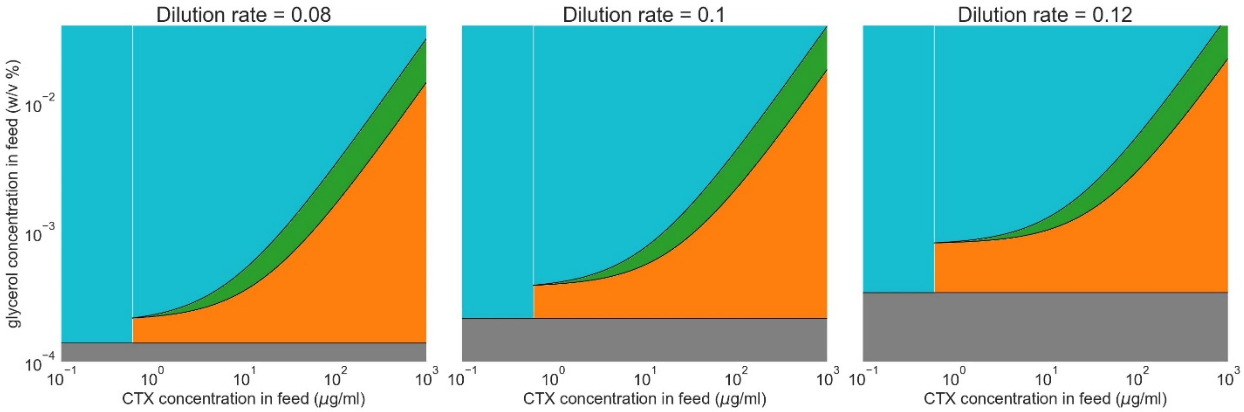

***Figure S5.*** *Predicted coexistence of TEM-52 and TEM-19 in the chemostat. Depending on the antibiotics in the feed, TEM-52 prevails (orange area), TEM-19 prevails (blue area) or the strains coexist (green area). High antibiotics favour the more resistant type, and high glycerol concentrations lead to high biomass, which favours the more susceptible type. In between there is an area of coexistence. The grey area denotes no survival of either strain.*

#### S4.4 Effect of changing conditions on coexistence of genotypes

The mechanism we show above is not dependent on the specific parameter settings. For example, the expression of the beta-lactamase could differ due to a different promotor, affecting the maximal degradation rate of both strains, or the cost of a different plasmid could incur additional or decreased costs in terms of growth rate to both genotypes. In Figure S6 we show that such changes only quantitatively affect our results: the antibiotic concentrations and cell densities at which coexistence are reached are slightly different, but the qualitative results remain the same. Note that we have shown quite large changes here, with changes in the growth rate that are near 20% of the base growth rate.

Although in a chemostat setting we do model dilution of the antibiotic to some extent (with the dilution rate of the chemostat), we looked at the effect of additional degradation (which could reflect pharmacokinetic effects). Even with an extra degradation term of 10 times the dilution rate, we see the same qualitative results (compare Figure S7 and Figure 4B in the main text).

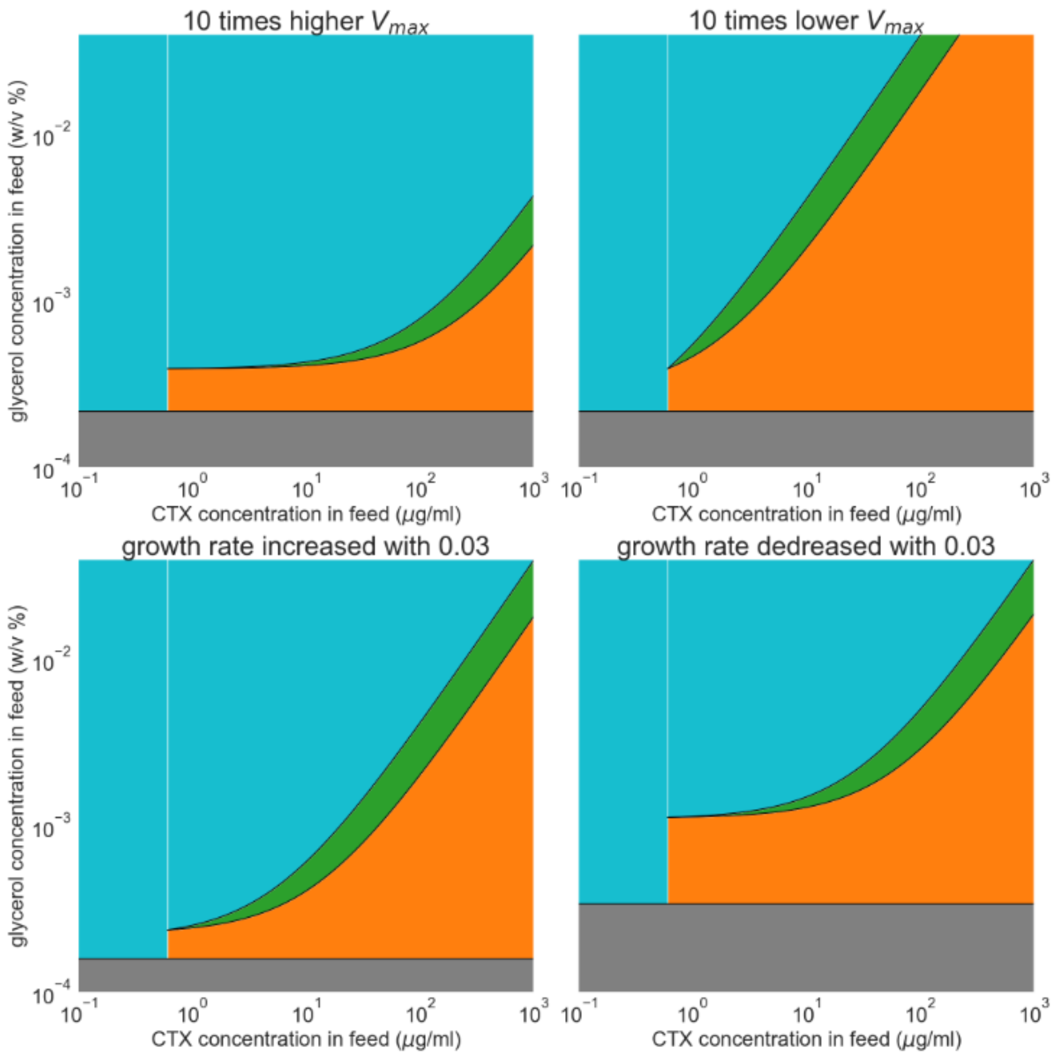

***Figure S6.*** *The effect of changes in parameters on the predicted coexistence of TEM-52 and TEM-19 in the chemostat. Depending on the antibiotics in the feed, TEM-52 prevails (orange area), TEM-19 prevails (blue area) or the strains coexist (green area).*

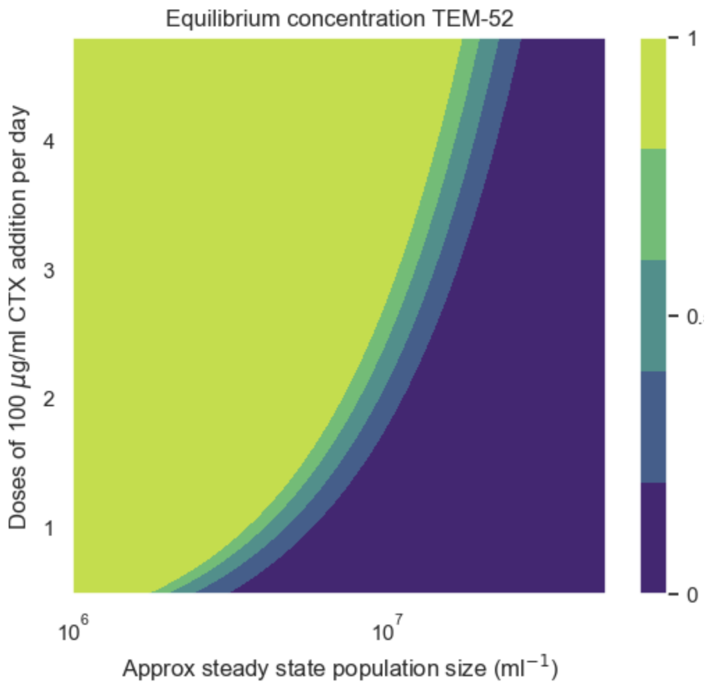

***Figure S7.*** *The effect of additional antibiotic degradation of 50 times the dilution rate on the predicted coexistence of TEM-52 and TEM-19 in the chemostat.*
